# Transforming Hyperspectral Data into Insight: The DREAM Approach for Pathology

**DOI:** 10.1101/2025.09.16.676582

**Authors:** Mason Hong, Daniel E.S. Koo, Jason Junge, Scott E. Fraser, Francesco Cutrale

## Abstract

Quantitative pathology remains limited due to the need for chemical staining and subjective interpretation of tissue features. Autofluorescence imaging offers a label-free alternative: however, high-dimensional excitation-emission datasets pose challenges for visualization and reproducible analysis. Here, we present Dimensionality Reduction for Enhanced Autofluorescence Microscopy (DREAM), a method that condenses multi-excitation emission spectra into a compact, information-rich format using phasor-based tools. Applied to unstained esophageal tissue samples, DREAM enables high-contrast visualizations that distinguish key histological structures without the need for exogenous labeling. Quantitative assessments across multiple datasets show DREAM improves colorfulness, sharpness, and consistency over single-laser acquisitions, supporting its potential to advance objective, label-free diagnostics through enhanced spectral visualization.

## 1. Introduction

Pathology plays a pivotal role in medical diagnostics by offering insights into conditions ranging from cancer to chronic diseases. Central to this discipline is the analysis of chemically stained tissue biopsies, commonly after Hematoxylin and Eosin (H&E) staining, to visualize tissue structures and cellular components. This process, however, is labor-intensive, time-consuming and subjective, as it requires expert pathologists to draw upon visual cues to assess the stained tissue slides under white light microscopy[1]. Pathologists evaluate these images by analyzing specific spatial features in the H&E-stained cells, including nuclear characteristics, cell elongation, cellular differentiation, tissue architecture, and overall tissue organization [2]. Cancerous cells often exhibit morphological abnormalities, such as irregular shapes, multiple large nucleoli, and reduced adhesion to neighboring cells, leading to disorganized tissue structures owing to their loss of specialized functions associated with their tissue of origin[3]. The collective interpretation of these features forms the foundation for determining the presence of malignancies within a sample. Despite the established diagnostic criteria, the evaluation process is adversely influenced by multiple inherent limitations, including inter-observer variability, reliance on subjective assessment and extended sample preparation procedures that can delay critical clinical decisions[4] .

Recent advances in label-free imaging techniques aim to address these limitations by leveraging intrinsic tissue properties for real-time, quantitative pathology. Label-free approaches bypass the need for chemical staining, instead relying on various optical methods to differentiate tissue types based on inherent physical and biochemical characteristics. Methods such as Stimulated Raman Scattering (SRS) and Coherent Anti-Stokes Raman Scattering (CARS) have been used to generate label-free tissue images by detecting vibrational signatures of biomolecules, offering contrast comparable to H&E staining[5, 6]. These techniques have been applied in glioblastoma tissue classification[7, 8] and metabolic mapping in cancerous and normal liver tissue [5]. Reflectance confocal microscopy allows for non-invasive, volumetric imaging of tissue microstructures [9-12], enabling real-time intraoperative assessments. Similarly, third-harmonic generation microscopy has demonstrated utility in imaging unstained human tissues, producing contrast-rich images based on structural properties rather than chemical stains[13-15].

Among the multitude of label-free methods, autofluorescence-based imaging has gained traction as a practical alternative due to its ability to exploit endogenous fluorescence signals emitted by biological molecules such as NADH, retinoids, and flavins. While much of autofluorescence research focuses on live samples, there are numerous contributors to the intrinsic signals of fixed tissues. Specific cell types and cellular components, such as membranes, cytoplasm, as well as components of the extracellular matrix, like collagen and connective tissues, will exhibit autofluorescence with specific excitation and emission spectral profiles [16, 17]. Metabolic biomarkers such as NADH in its free and bound form, are known to emit autofluorescence[18], permitting metabolic imaging in multiple settings, such as in cell cultures, pancreatic cancer and breast cancer both in live imaging[19-21], and even after fixation[22]. Autofluorescence imaging has been utilized to analyze metabolic disruptions, including those in photoreceptors [23, 24], and to enhance histopathology through spectral characterization of intrinsic fluorophores[16]. The integration of spectral detection with autofluorescence expands the scope of information that can be extracted, enabling precise tissue classification based on spectral fingerprints. For example, hyperspectral autofluorescence has been used to identify amyloid deposits in neurological disorders[25], and to distinguish cancerous from non-cancerous tissues based on metabolic differences[20, 26].

Building on these spectral fluorescence advancements, we introduce Dimensionality Reduction for Enhanced Autofluorescence Microscopy (DREAM), a powerful approach that synthesizes spectral information from multiple excitation wavelengths into a single high-information-content fluorescence emission spectrum. This method enhances spectral contrast and improves the differentiation of tissue types and pathologies. DREAM-generated spectral data are analyzed using Spectrally Encoded Enhanced Representations (SEER)[27], which employs phasor analysis to convert spectral information into quantitative, interpretable color maps. By leveraging hyperspectral autofluorescence and phasor-based analysis, we demonstrate that DREAM provides an objective, quantitative alternative to conventional pathology, reducing diagnostic time while enhancing contrast and specificity in clinical assessments.

## 2. Methods

### 2.1 Sample Preparation and Hematoxylin and Eosin (H&E) Imaging

Slides were prepared and stained at the Translational Pathology Core of the University of Southern California (USC). Each sample set included two vertically sequential 5μm sections (Fig. 1a). One section was left unstained for autofluorescence imaging (Fig. 1b); the other was processed utilizing standard protocols and stained with H&E (Fig. 1c). This sequential section approach provides a reasonable comparison for stained and unstained samples, with a minimal tissue offset of one section thickness (5μm). Stained slides were imaged at the Translational Pathology Core of USC following established protocols, utilizing a Hamamatsu Nanozoomer S60 digital slide scanner, acquiring images in bright field mode at 20× magnification. H&E pathology images were scored by registered pathologists, identifying areas with lesions (Fig. 1g). 19 datasets were acquired at different position from three patient-derived specimens with metaplasia assessment.

**Figure 1:**
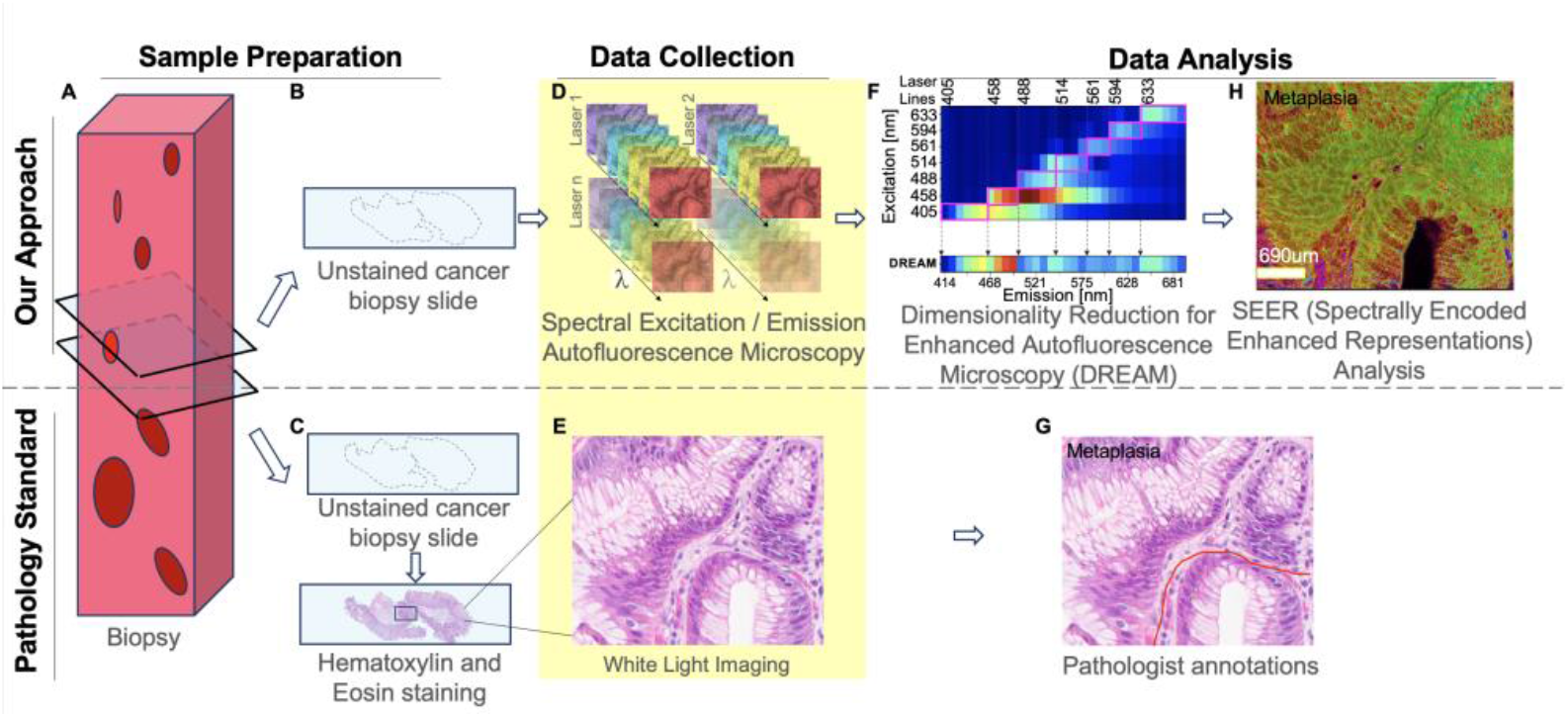
Approach overview to compare DREAM to conventional pathology. (A) Two vertically sequential sections are taken from a sample. (B) One section is left untreated for label-free hyperspectral autofluorescence analysis while (C) the other sample is conventionally processed and H&E stained. (D, E) The untreated section is imaged using hyperspectral fluorescence microscopy; the H&E section is imaged using standard white light microscopy. (F) The hyperspectral data of the untreated slide is processed using Dimensionality Reduction for Enhanced Autofluorescence Microscopy (DREAM) and analyzed using Spectrally Encoded Enhanced Representations (SEER; here gradient descent, harmonic = 1), to reproducibly and automatically highlight spectrally distinct elements. (G) the H&E images are reviewed and annotated by a pathologist.

### 2.2 Spectral Excitation and Emission Autofluorescence Imaging and Maps

Unstained cancer slides were imaged using a confocal laser scanning microscope (Zeiss LSM880 inverted, Jena, Germany) set to operate in lambda mode with Quasar 32-channel detection (wavelength range of 410–692nm;8.8nm spectral bands), utilizing single-photon excitation. Samples, mounted on an imaging slide, were imaged at room temperature. Pathologists’ annotations were used as a guide for identifying areas of focus for imaging. Autofluorescence spectral imaging datasets were acquired on a single plane in 2D and tiled mode, to permit the analysis of samples larger than the microscope’s field of view (FOV). Each tiled dataset was positioned to include both cancerous and non-cancerous tissue regions.

For each analyzed region, we collected seven datasets, each excited with a different laser wavelength (405nm, 458nm, 488nm, 514nm, 561nm, 594nm, 633nm), utilizing the full detection range (410nm–692nm, 32 channels), with appropriate laser-blocking filters, specifically MBS-405 for 405nm ex. MBS 458 for 458nm ex, MBS 488 for 488nm ex, MBS 458/514/594 for 514nm ex, MBS 488/561 for 561nm ex, MBS 488/594 for 594nm ex, MBS 488/633 for 633nm ex. Single-band filters were used where possible to maximize the collected autofluorescence emission signal. Datasets were acquired using a 63× oil immersion lens (Plan-Apochromat 63x/1.40 Oil DIC, Zeiss, Jena). Microscope settings were optimized to fill 80% of the dynamic range in detection to avoid saturation. Pixel size (0.094um), pixel dwell time (3.15us), gain (700 for 405 excitation, 780 for all others) and laser power (0.2% for 405nm, 1.5% for 458nm, 0.2% for 488nm, 0.5% for 514nm, 0.3% for 561nm, 8% for 594nm, 22% for 633nm) were maintained consistently for each illumination line within experiments. The output for each of the 10 slides, was a 4-dimensional dataset with dimensions (x, y, excitation wavelength, emission wavelength), where x and y are the pixel positions in the image, excitation wavelength corresponds to one of the seven laser excitations, and emission wavelength represents the 32 spectrally resolved detection channels (Fig. 1d).

The inherently 4-dimensional datasets obtained from the confocal microscope are represented as excitation-emission maps, which visualize fluorescence emission intensity as a function of excitation wavelengths (Fig. 1f, Fig. 2). These maps provide insight into the spectral composition of tissues, with colors representing intensities, scaled relative to the maximum value within each sample, to preserve the relative intensity ratios of the multiple fluorophore contributions.

**Figure 2.**
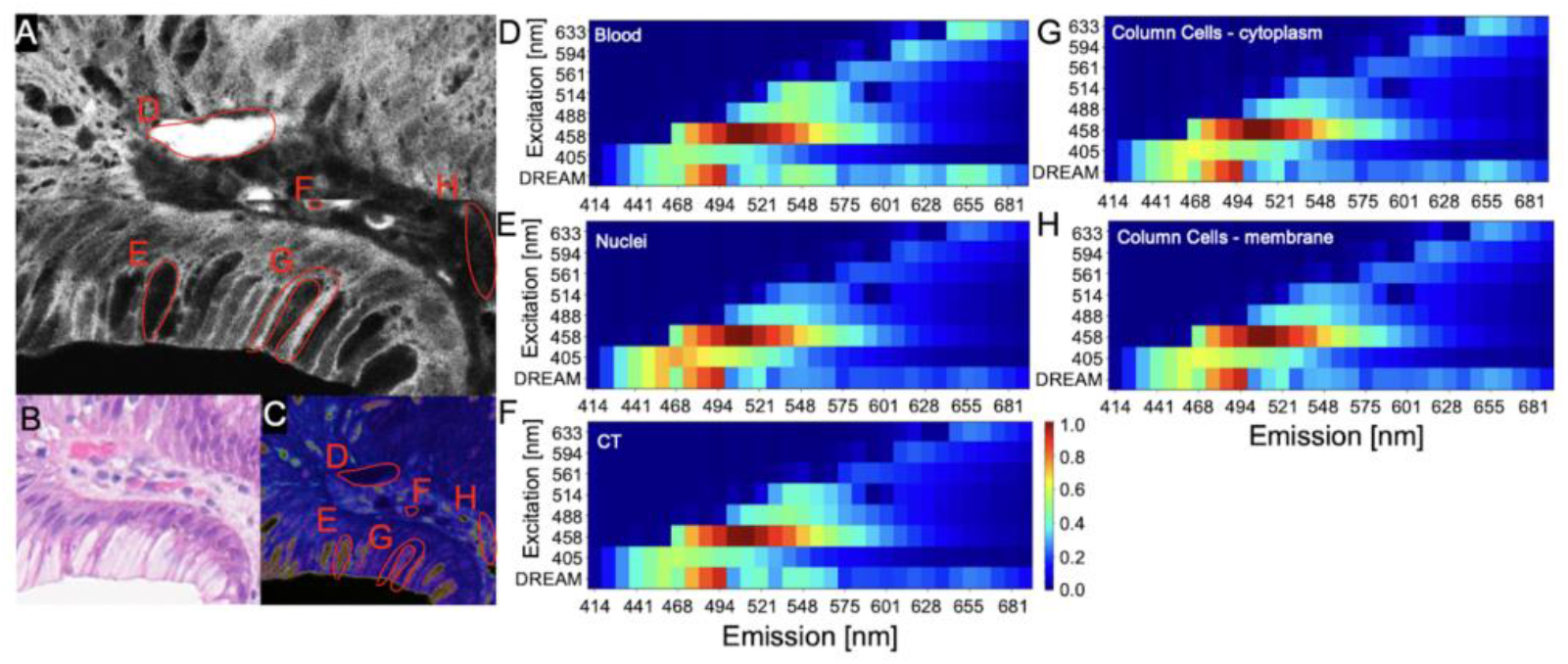
Spectral fingerprinting of different biological features. (A) Average spectral intensity grayscale image of a single z-slice of an unstained pathological tissue containing the five different types of cellular components. (B) H&E stained image of the same region as shown in A. (C) SEER image of the same region as shown in A at 458nm excitation. Normalized heatmaps representing the emission profiles of (D) Blood, (E) Column Cells - cytoplasm, (F) Nuclei, (G) Column Cells - membrane, (H) Connective Tissue for both DREAM and all excitation wavelengths (405 to 633nm).

We identified distinct structural and cellular features within the pathology slides (Fig. 2a-c) and reported their excitation-emission maps (Fig. 2d,h). These maps reflect the combination of different endogenous fluorophores within the features, responding to varying excitation wavelengths. The differences observed across tissue structure maps arise from the diverse molecular composition and varying metabolic states, potentially facilitating spectral differentiation between key tissue components such as nuclei, cytoplasm, membranes, and extracellular matrix.

### 2.3 Dimensionality Reduction for Enhanced Autofluorescence Microscopy (DREAM)

To simplify the performance and interpretation of hyperspectral autofluorescence imaging, we introduce DREAM, a method that synthesizes the four-dimensional (x,y,excitation, emission) spectral information into a single, high-information-content three-dimensional (x,y, emission) fluorescence dataset. Each 4-D excitation-emission dataset contains fluorescence emission spectra acquired following individual excitations by seven distinct wavelengths (here 405, 458, 488, 514, 561, 594 and 633nm), with each excitation generating a 32-channel emission spectrum. In DREAM, we construct a single composite emission spectrum by selecting the emission channels that follow each laser excitation, starting from the excitation channel itself and including all subsequent emission channels up to, but not including, the excitation channel of the next laser in the sequence (Fig. 1f). For example, after 405 nm excitation, we retain the emission channels up to the start of the 458 nm dataset, repeating the procedure for the latter laser line. This strategy ensures continuous spectral coverage without gaps, while avoiding cross-contamination between excitations. (Fig. 1).

This approach is motivated by the challenge of incomplete sampling when relying on any single excitation wavelength. Biological tissues contain a complex mix of autofluorescent biomarkers, such as NADH, FAD, retinoids, collagen, and porphyrins, each optimally excited by a different laser and emitting at unique spectral ranges. As such, single-laser acquisitions provide only a partial representation of the tissue’s biochemical makeup; for instance, a fluorophore excited at 633 nm may emit no detectable signal under 405 nm illumination. DREAM addresses this by integrating the most information-rich emission segments from each excitation channel, allowing for a broader and more holistic capture of spectral content. By focusing on the high-yield spectral windows while discarding low-information tails, DREAM enhances sensitivity and spectral contrast (Fig. 3) while streamlining the dataset by excluding redundant spectral content and reinforcing excitation-specific details.

**Figure 3.**
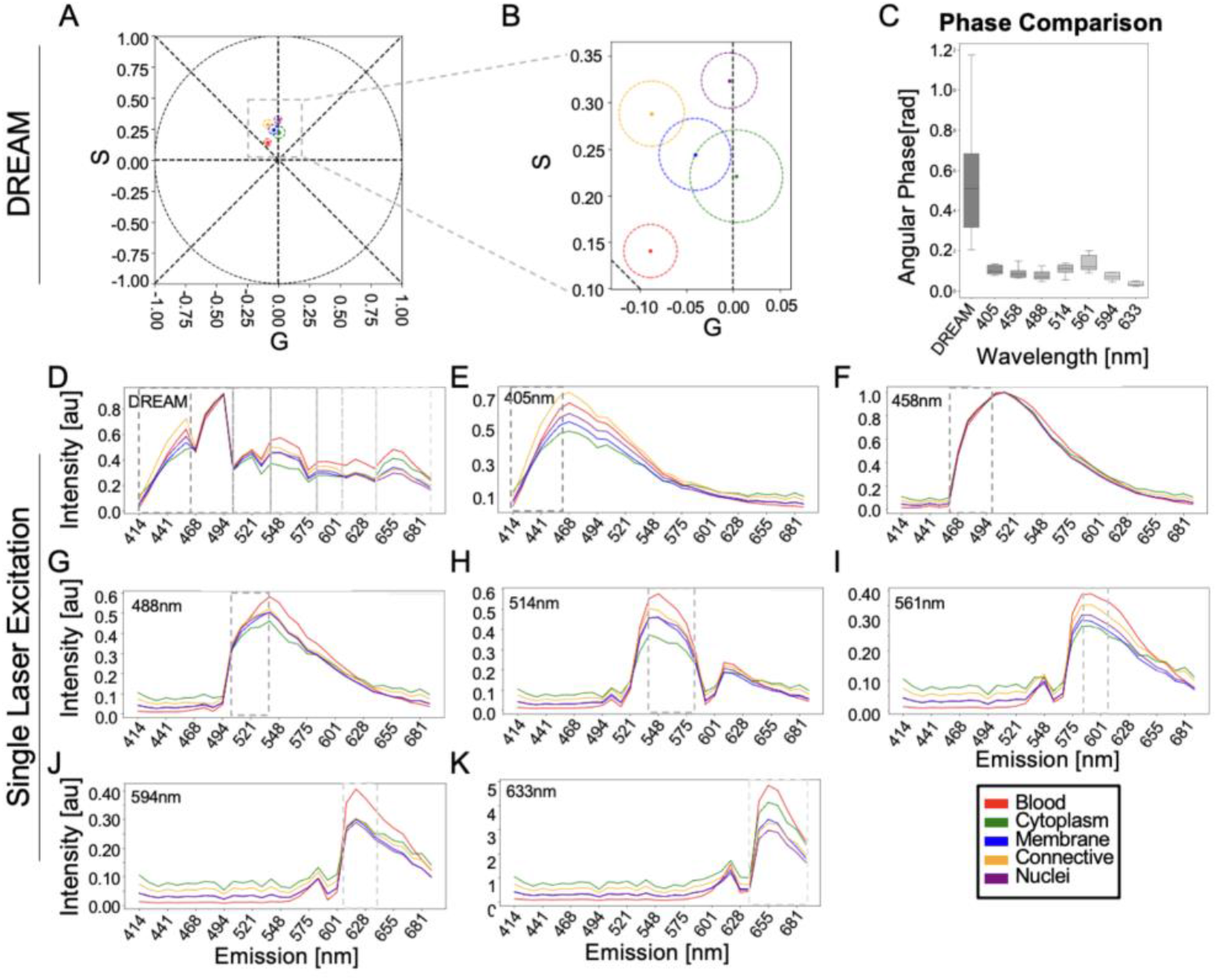
Spectral comparison of DREAM and single-excitation autofluorescence datasets. Extension of the feature-level excitation/emission mapping shown in Fig. 2 by comparing the spectral separability of DREAM with that of conventional single-laser acquisition. (A) Phasor plot of DREAM-processed data showing the distribution of pixels corresponding to five biologically distinct tissue features: blood, cytoplasm, membrane, connective tissue, and nuclei. (B) Zoomed-in view of the phasor space highlighting the 5% closest pixels to the mean phasor coordinate for each feature, visualized as colored circles with radius equal to the maximum distance among those pixels. These radii represent the smallest phasor distance enclosing the 5% nearest-to-mean phasor subset pixels for each tissue type. (C) Angular Phase spread across datasets, reported as boxplots, defined as the difference between the smallest and largest phasor angle among the five features. The center line indicates the median, box edges represent the interquartile range (IQR, 25th–75th percentiles), and whiskers extend to 1.5× the IQR. DREAM shows a larger median phase spread than any single-excitation image, indicating greater separation of tissue spectra in the phasor domain. (D) Average emission spectra from the DREAM dataset for each feature reveal distinct spectral fingerprints. Dashed boxes, in grayscale, indicate the spectral bands of relevance for the DREAM approach. (E–K) Emission spectra of the same tissue features from each individual excitation wavelength (405, 458, 488, 514, 561, 594, and 633 nm) across 19 datasets acquired from 3 different tissue samples highlight DREAM’s increased distinction in phasor spectral signatures across all tissue types. The bottom-right legend applies to all panels in this figure.

The resulting DREAM-generated dataset is a dimensionality-reduced but biologically comprehensive view of tissue autofluorescence. This compact representation is particularly well-suited for downstream phasor-based spectral analysis [27-29], to enhance visualization and differentiation of subtle differences across tissue structures. By condensing the most information-rich signals into a single, optimized spectrum, DREAM enhances the sensitivity and contrast of images from intrinsic fluorescence (Fig. 3). The spectral window selection used in this work is rule-based and can hence be expanded, for instance by adding a long-pass segment, to capture large Stokes-shit emitters or to support other dimensionality-reduction workflows, all while preserving the original channel intensities. The enhanced contrast and spectral sensitivity observed with DREAM, provides a more informative and streamlined analysis of complex hyperspectral autofluorescence imaging data, highlighting its utility for label-free histopathology and paving the way for future extensions to more complex unmixing and classification tasks.

### 2.4 Phasors and SEER

DREAM-processed spectra are visualized using Spectrally Encoded Enhanced Representations (SEER)[27], which applies spectral phasor analysis to transform three dimensional hyperspectral data, comprising spatial (x,y) and wavelength (λ) dimensions, into quantitatively interpretable RGB images. SEER uses phasor analysis for fluorescence microscopy[29-31], which transforms the emission spectrum at each pixel into its real and imaginary Fourier components. These components define a coordinate in a two-dimensional space known as the phasor space, where each point (g,s) represents one autofluorescent spectrum. The phasor plot is represented as a 2D histogram where each bin contains a collection of pixels whose spectra share similar characteristics based on features such as shape, intensity and peak position. Spectra with similar shapes map to nearby phasor coordinates, and contiguous high-density regions, we define as clusters, correspond to recurring spectral populations [32]. The clusters of the phasor plot histogram capture the population-level distributions of pixels in the dataset from a spectral perspective [29]. SEER then applies tailored, distribution-adaptive color maps to assign colors to specific bins on the phasor plot, generating RGB images where similar colors indicate corresponding spectral signatures. This results in a continuous and adaptive visualization where pixels with distinct spectra appear in contrasting colors, effectively differentiating fluorophores based on their spectral properties [27].

SEER visualized images of DREAM datasets (DREAM-SEER images) significantly enhance the visual contrast of intrinsic autofluorescent signals, enabling distinction of tissues and cells solely based on their spectral characteristics outperforming PCA and ICA in photon-starved fluorescence settings [27]. Spectra represented on the phasor plot will be distributed according to a phasor angle, related to the spectral phase or weighted center of the spectrum, and a phasor radius, proportional to the overall intensity of signal, with lower-signal spectra closer to the central position of the phasor. In this work we utilize the phasor gradient descent color map, a reference palette designed to emphasize spectral variations in low-intensity signals, typically overlooked in other color re-mappings. This map highlights subtle spectral differences even in dim autofluorescent signals, enhancing their visual detectability. The color contrasts observed in DREAM-SEER images reflect not only large shifts in emission spectra but also lower signal-to-noise, spectrally distinct populations that might overwise be masked by more intense signals [27]. In the pathology samples in this work, the color coding reveals discrete tissue features, such as nuclei, blood and connective tissue, based on their composite spectral properties, which arise from differing combinations of endogenous biomarkers.

### 2.5 Image processing

Before undergoing DREAM-SEER analysis, autofluorescence spectral images were pre-processed using FIJI [33] to segment different features for phasor analysis and for quantification of the perceptual image quality in all SEER images (Fig. 2). Referencing the corresponding H&E images, we created Regions of Interest (ROIs) around selected biological features such as nuclei, column cells, blood cells and connective tissues, on a single, high intensity channel within the autofluorescence spectral datasets in FIJI by utilizing the polygon selection function. We then converted the ROIs to a mask by using the Edit-Clear function in FIJI. Finally, the mask was applied to all 32 channels.

DREAM analysis was performed using custom python scripts. Briefly, for each spectral excitation/emission datasets, the excitation wavelength was associated with a known acquisition channel in the 32-channel detector. To build the DREAM dataset, we identified the emission channels that followed each excitation wavelength, starting from the excitation channel itself and continuing through the subsequent channels, stopping before the next excitation wavelength’s channel (Fig. 1f). For instance, for the 405nm excitation, the dream selection includes channels from 405nm through to, but not including, the 458nm excitation channel. This same approach was applied across all laser lines. The resulting non-overlapping segments from all excitations concatenated to form a single, continuous 32-channel emission spectrum per pixel. This deterministic rule avoids synthetic spectral overlap between excitation datasets and maintains linearity for downstream analysis. It is well suited for short-to-moderate Stokes-shift autofluorescence and can be extended, by adding a long-pass segment, to fluorophores with large Stoke-shifts. Although DREAM is a compression, the retained spectra (Fig. 2 are sufficient to support high-contrast SEER visualization, as evidenced by greater phasor Angular Phase separation (Fig. 3) and significantly higher image quality metrics relative to single-excitation images (Figs. 4, 5, 6).

**Figure 4.**
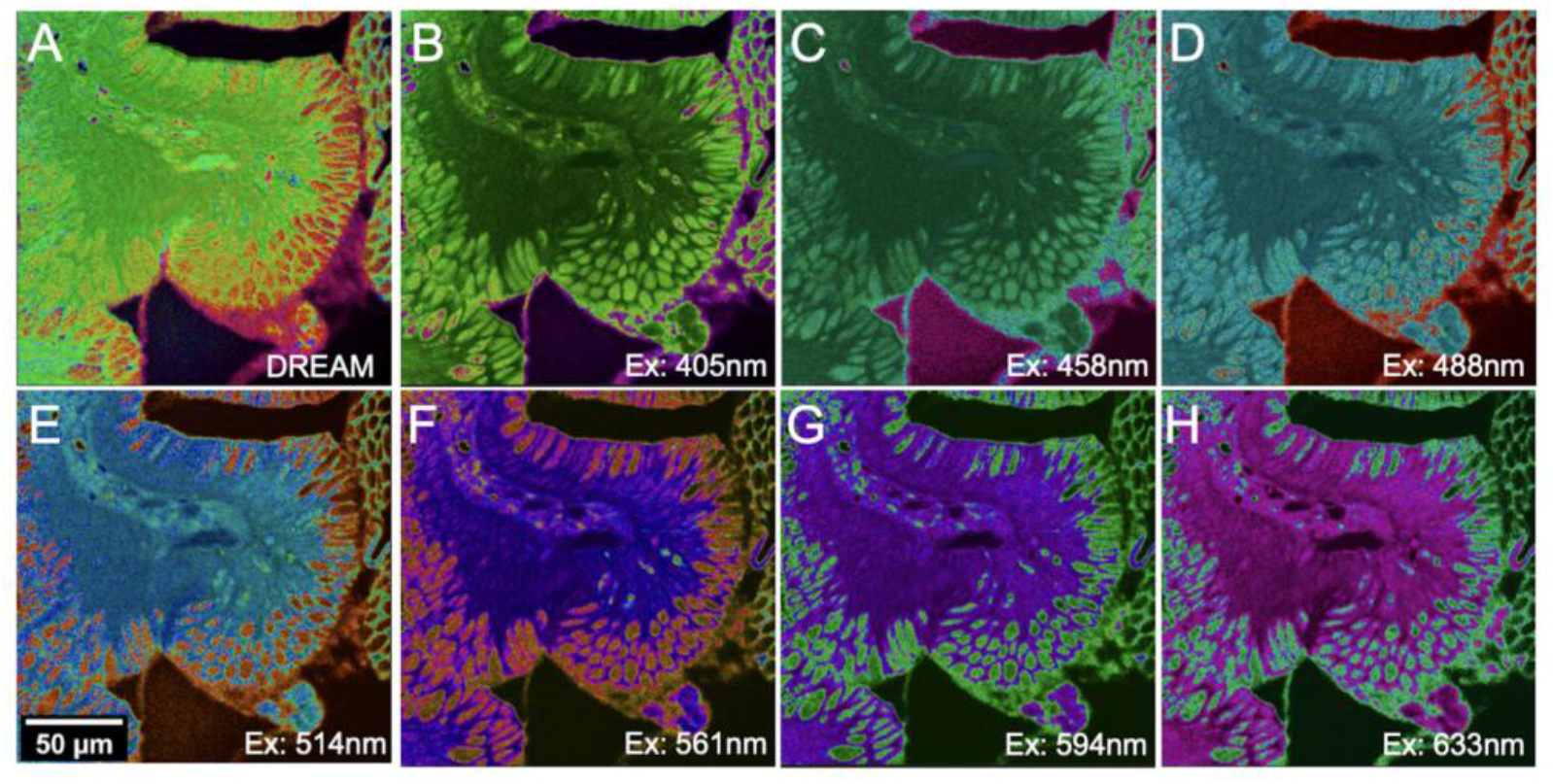
SEER visualizations of DREAM and single-excitation autofluorescence datasets. SEER-rendered images for the same tissue region acquired using DREAM (A) and individual laser excitations: 405 nm (B), 458 nm (C), 488 nm (D), 514 nm (E), 561 nm (F), 594 nm (G), and 633 nm (H). All images use the same gradient descent SEER color map, applied independently to each dataset’s phasor distribution. The DREAM image (A) shows enhanced contrast and differentiation between the tissue features of columnar cells, nuclei, blood, and connective tissue, relative to single-laser acquisitions. Single-excitation datasets appear to highlight specific components, such as cytoplasm in 405 nm and membranes in 561 nm, but DREAM provides the most comprehensive spectral encoding, revealing finer distinctions and broader spatial detail.

**Figure 5.**
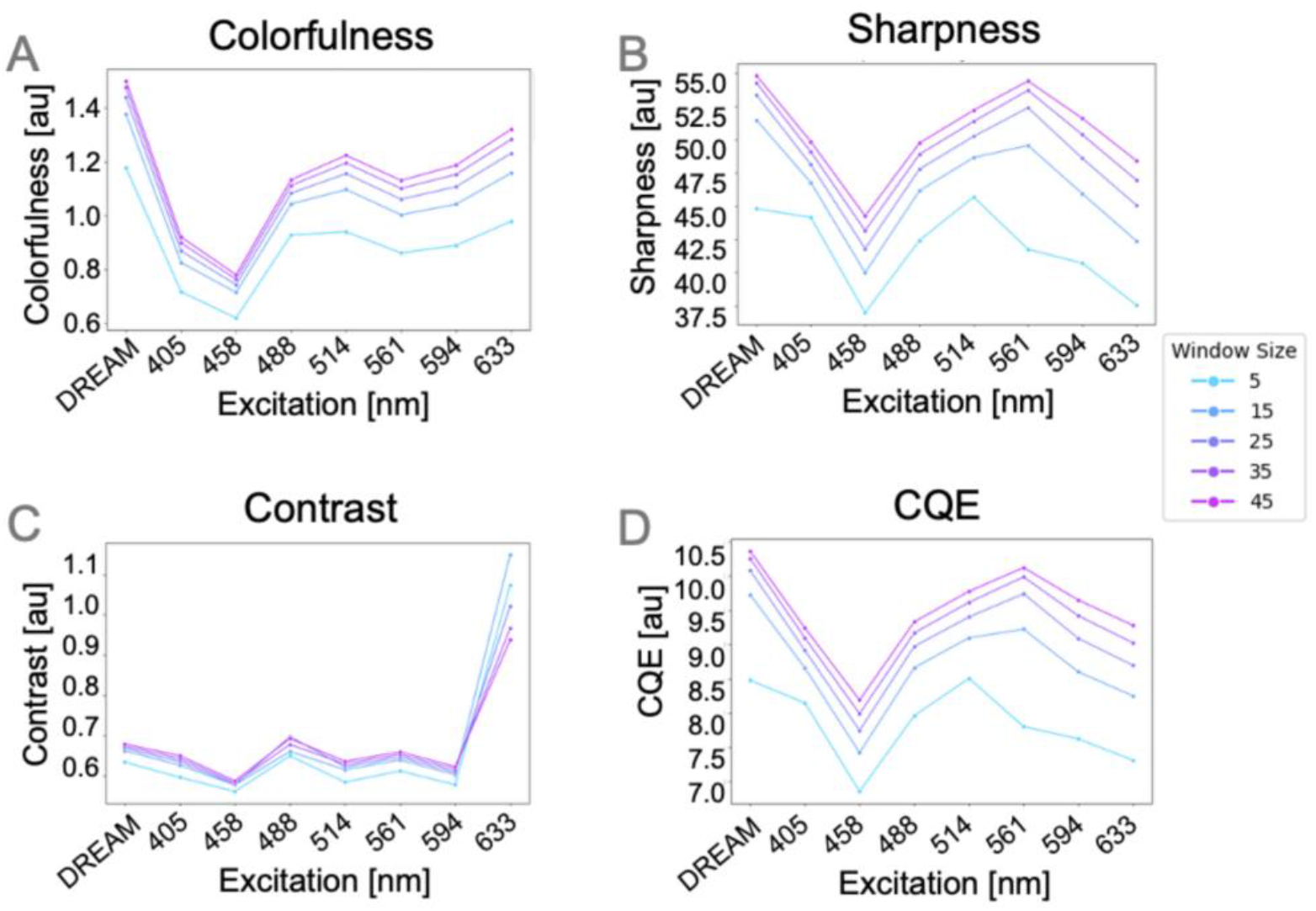
Quantitative evaluation of SEER image quality across spatial scales. Perceptual image quality metrics (A) colorfulness, (B) sharpness, (C) contrast (CRME), and (D) Color Quality Enhancement (CQE) computed from SEER images shown in Fig. 4 for DREAM and single-laser excitation datasets. Each curve represents the metric value computed using a sliding window of size 5, 15, 25, 35, or 45 pixels, corresponding to different biological feature scales. DREAM consistently achieves the highest colorfulness and sharpness across all window sizes. While the 633 nm dataset shows higher localized contrast, DREAM maintains the highest overall CQE scores, highlighting its superior perceptual image quality over single-wavelength excitations.

**Figure 6.**
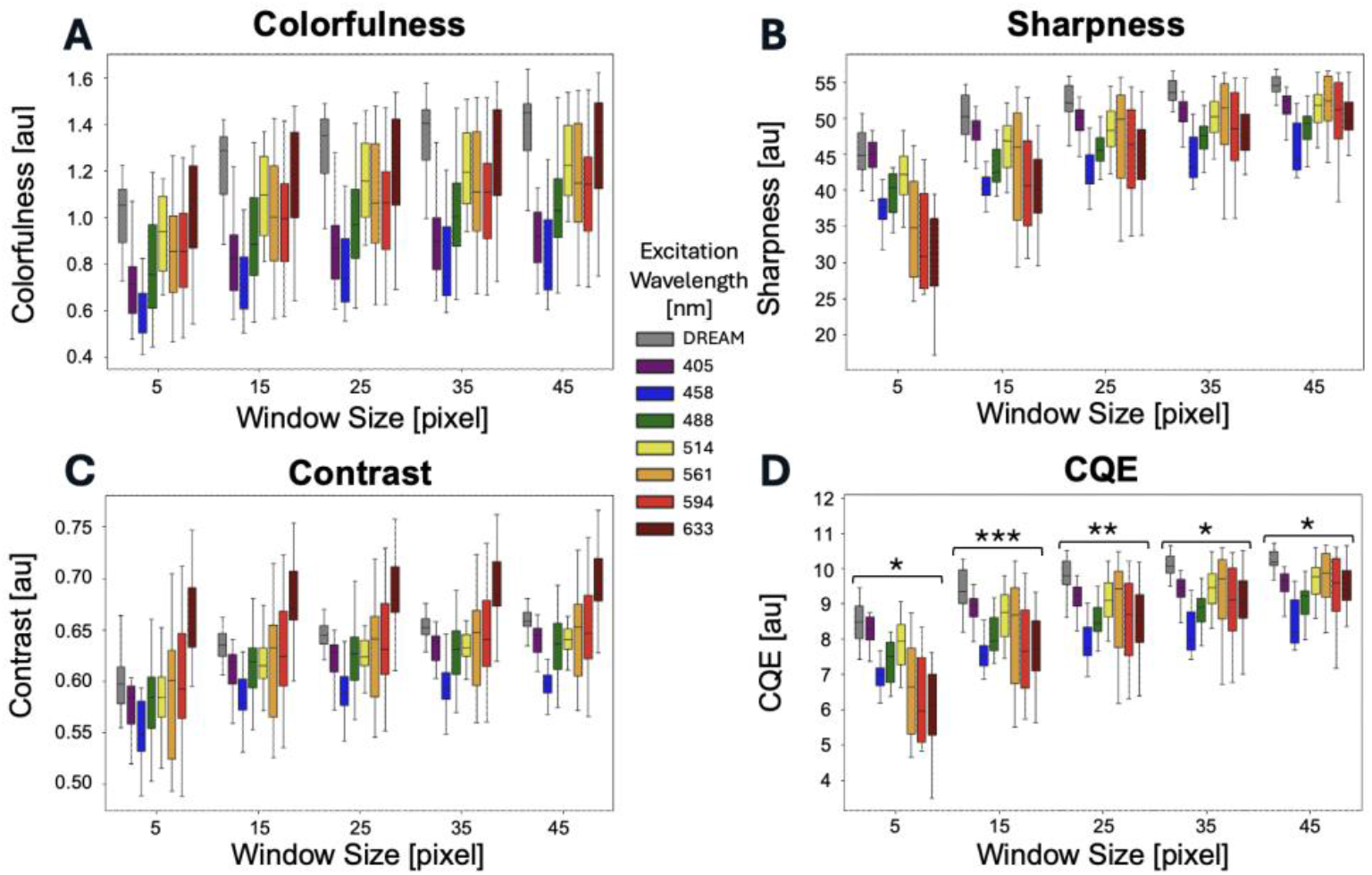
Aggregated image quality metrics across 19 SEER datasets from three samples. Box plots summarize (A) colorfulness, (B) sharpness, (C) contrast (CRME), and (D) Color Quality Enhancement (CQE) metrics calculated from 19 SEER-processed datasets acquired from three distinct tissue samples from different patients. Metrics were evaluated over five sliding window sizes (5–45 pixels) to capture variations across spatial scales. Each box plot represents the distribution of metric values across datasets for a given excitation wavelength. The center line indicates the median, box edges represent the interquartile range (IQR, 25th–75th percentiles), and whiskers extend to 1.5× the IQR. The DREAM method (gray) consistently achieves the highest median colorfulness, sharpness, and CQE across all window sizes. While the 633 nm excitation yields the highest contrast, it also shows wider variance. DREAM, by contrast, demonstrates more compact distributions, especially for CQE and sharpness at larger window sizes, suggesting enhanced robustness and consistency in image quality across varying tissue features. *P value<0.05, **P value<0.01, ***P value<0.001.

SEER analysis was performed using HySP v0.9.20 software[29] (www.bioimage.usc.edu/software.html) both on MacOS (13.6-14.2.1) (2.3 GHz Quad-Core i5, 8 GB ram, Intel Iris Plus Graphics 655 1536 MB) and a Windows PC equipped with (Intel(R) Core(TM) i7-7820X CPU - 3.60GHz, 128 GB,NVIDIA GeForce GTX 1060 6GB).

The phasor coordinates g and s for a given pixel (x, y, z, t) at harmonic h = 1 are defined by the following equations:

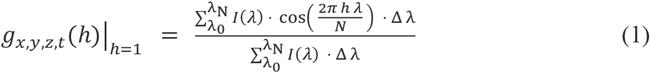

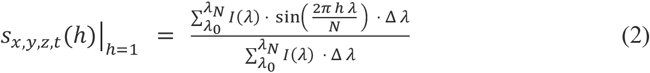

Where *I*(*λ*) is the fluorescence intensity at emission wavelength *λ*; *λ*_0_ and *λ*_*N*_ are the lower and upper bounds of the emission wavelength range; Δ*λ* is the spectral step or channel resolution (for our system 8.8 nm); N is the total number of spectral channels; k is the harmonic used for phasor transformation (k=1 for this work); g and s are the real (cosine) and imaginary (sine) components of the phasor, respectively. The transformation yields a phasor plot where each point represents a unique emission spectrum. Spectra with similar shapes cluster together, allowing complex datasets to be visualized and interpreted using color-encoded SEER maps.

Prior to SEER and phasor analysis, all datasets were thresholded. Threshold values were selected using FIJI’s auto-contrast algorithm, adapted from the Java code, (https://github.com/imagej/ImageJ/blob/706f894269622a4be04053d1f7e1424094ecc735/ij/plugin/frame/ContrastAdjuster.java#L780). The selected values from the auto-contrast algorithm were multiplied by 3 (value obtained empirically) to produce threshold values.

We applied three sets of phasor denoising, via median filter of the image real and imaginary coefficients[29]. Datasets are visualized using SEER’s gradient descent original map with the autoscale submode, highlighting the spectral differences between low-intensity spectral signals[27]. To quantify the separation of signals in phasor space, we utilized Angular Phase difference. For each dataset, we computed the mean phasor coordinate 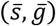 for each annotated tissue ROI after thresholding and denoising, converted it to a phase angle

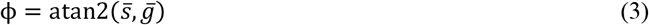

in radians, ensuring angles were between [0,2*π*], and then defined the dataset’s angular phase difference as

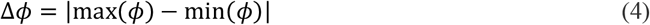

To minimize the contribution of laser reflection artifacts, a masking step was applied to the 633 nm datasets and propagated to the DREAM datasets for all samples’ datasets. Masking was performed by examining channel 24 of the 633 nm dataset, which corresponds to the emission band immediately preceding the laser blocking filter. Pixels with high intensity values, typically above 15,000-17,000 digital levels, were identified as likely laser reflection artifacts. These pixels were cross-referenced with spectral clusters on the phasor plot in HySP, allowing us to localize the region of interest (ROI) corresponding to the laser scattering. Using the ROI selection tool in HySP and some post-processing, a binary mask was generated from the 633 nm dataset and applied to both the DREAM and 633 nm datasets to suppress these artifacts. The masked datasets were then exported as Tiff File stacks for subsequent analysis. This procedure affected on average 0.23% +/-0.18% of pixels per image and targeted only localized hotspots; analyses and perceptual metrics were computed on the remaining pixels. No comparable artifacts were detected for the other excitation wavelengths using the same screening. Given the small affected area and the removal of non-fluorescent scatter, this masking step is not expected to compromise spectral integrity or interpretability of the fluorescence data.

During phasor transformation and SEER visualization, the masking process occasionally introduced disconnected spectral bins, accounting for accounting for 0.045 % ± 0.015 % of the pixels, on the phasor plot that disrupted SEER color mapping. To address this, we used the *connected_regions* function from SciPy to identify the largest contiguous region in phasor space, containing the majority of pixels and spectral diversity. All bins not connected to this main region were removed prior to SEER map application, preventing a negligible set of outliers from influencing palette endpoints.

### 2.6 Quantification of perceptual image quality

To evaluate the image quality offered by DREAM-SEER compared to conventional single-excitation SEER, we utilize an established method for quantifying perceptual image quality[34] based on three key perceptual metrics sharpness, contrast, colorfulness, and a single summary metric, the Color Quality Enhancement (CQE) score, which integrates them. These no-reference metrics are used to assess visual clarity, texture differentiation, and chromatic richness in image data.

Sharpness refers to the preservation of fine details and edges, and in our approach, it is quantified following the method defined by Gao et al. [34]. The Sobel edge detection algorithm is first applied to each RGB component, generating binary edge maps. These maps are multiplied with the original image values to produce grayscale edge-enhanced images. A Weber contrast-based enhancement metric, known as the Measure of Enhancement (EME), is then applied over 3×3 overlapping windows on each grayscale edge image. The final sharpness score is a weighted combination across color channels:

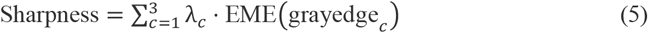

with:

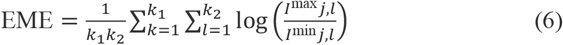

where *I*^max^*j, l and I*^min^*j, l* represent the maximum and minimum intensities within the local M by N pixels window at position (j,l), and *λ*_*c*_ are NTSC-based channel weights for red (0.299), green (0.587), and blue (0.114), while *k*_1_, *k*_2_ are the number of vertical and horizontal window positions, respectively. For overlapping windows with stride 1, *k*_1_ = *H* − *M* + 1 and *k*_2_ = *W* − *N* + 1, where H and W are the image height and width in pixels.

Contrast is measured using the logarithmic Michelson-based enhancement measure (AME), adapted for color images and designed to increase with perceived contrast:

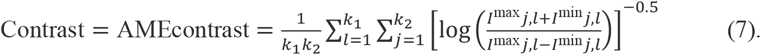

Where *k*_1_, *k*_2_, j, l have the same definition above. This formulation reverses the traditional AME output to ensure higher values correspond to higher visual contrast, aligning with human perception.

Colorfulness quantifies chromatic intensity and variation in the image and is calculated using previously reported logarithmic formulation [34]. This version provides a stronger correlation with human perception by leveraging opponent color channels and normalizing variance by the mean:

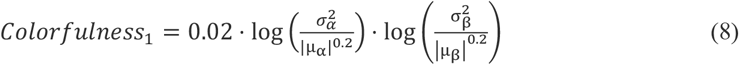

Here, *α* = *R* − *G* and β = 0.5 · (*R* + *G*) − *B* represent the opponent color axes, while μ and σ^2^ denote the mean and variance along each axis.

Color Quality Enhancement (CQE) is computed as a linear combination of the three perceptual metrics above. The CQE score provides a compact representation of overall image quality:

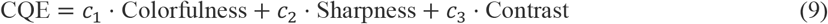

where *c*_1_, *c*_2_, *c*_3_ are weighting coefficients (e.g., for generic distortions: *c*_1_ = 0.2946, *c*_2_ = 0.3483, *c*_3_ = 0.3571). This framework enables objective, perceptual-quality comparison between DREAM-SEER and conventional SEER approaches across biological images.

To ensure spatial consistency and to account for local variations in image quality, each of the three metrics, sharpness, contrast and colorfulness, was computed over a rolling window of *n* × *n* pixels across the image with n=[5,15,25,35,45]. These windowed measurements were then averaged across the full image to yield a single representative value per metric. The CQE score was subsequently derived using the averaged values, preserving both global and local perceptual characteristics while enabling robust comparison. At each spatial window size, we compared CQE from the DREAM images against each single-excitation image (λ = 405–633 nm) using a paired Wilcoxon signed-rank test.

## 3. Results

### 3.1 Autofluorescence Excitation-Emission Fingerprinting of Tissue Features

To characterize autofluorescent signatures in pathological tissue, we constructed excitation-emission maps by segmenting biologically relevant features in unstained autofluorescence datasets. Fig. 2a shows the average fluorescence intensity image along the wavelength dimension, acquired under 488 nm excitation, where regions of interest were manually segmented based on correspondence with standard H&E (Fig. 2b). These include blood vessels, nuclei, connective tissue (CT), and columnar epithelial cells, with the latter further subdivided into cytoplasm and membrane domains. For illustration, we show the SEER visualization of the same field at 458 nm excitation in Fig. 2c. Each selected region is used to compute average excitation-emission maps, shown in Fig. 2d–2h, where the x-axis represents emission wavelength (410–692 nm) and the y-axis denotes excitation wavelength (405– 633 nm). Intensities are scaled relative to the maximum intensity per region, preserving relative brightness across excitation channels.

The resulting excitation-emission maps display distinctive spectral fingerprints for each tissue component. Blood cells show a strong emission peak in the 655 nm region under 633 nm excitation (Fig. 2d), while nuclei exhibit a more pronounced signal in the 488–521 nm range when excited at 405 nm (Fig. 2f). Connective tissue is characterized by weak emission under longer excitation wavelengths (561–633 nm) and a prominent response at 488 and 514 nm excitations with emission in the 521–575 nm band (Fig. 2h). Columnar cell cytoplasm and membrane show similar excitation-emission profiles (Figs. 2e and 2g), with only minimal differences, suggesting that their endogenous fluorophore content may be largely overlapping. These excitation–emission profiles demonstrate quantitative discriminability, with spectral parameters that distinctly characterize each tissue component. The resulting separability across spectral channels facilitates differentiation of blood, nuclei, connective tissue, and epithelial structures within the autofluorescence dataset.

### 3.2 DREAM Spectra Improve Feature Differentiation

While excitation-emission maps offer insight into molecular contrast, DREAM condenses this high-dimensional data into a single 32-channel emission spectrum by selecting the most informative emission range for each excitation wavelength. In Fig. 3, we evaluate whether the DREAM-processed spectra preserve—and potentially enhance—the spectral uniqueness of the tissue components identified in Fig. 2. Fig. 3a presents the phasor plot computed from DREAM spectra of blood, nuclei, connective tissue, cytoplasm, and membrane regions. The zoomed inset (Fig. 3b) shows that each feature occupies a distinct position in phasor space, indicating spectral separability. This distinction is further supported by their corresponding emission profiles in Fig. 3c, which illustrate the DREAM-processed spectra for each feature.

For comparison, Fig. 3d–3j display the emission spectra for each tissue feature under single excitation wavelengths (405–633 nm). Some individual wavelengths, such as 514 nm and 561 nm (Figs. 3g and 3h), yield moderately distinct spectral separations between features. However, other excitations (e.g., 405 nm, 458 nm, and 594 nm) provide limited discrimination. Notably, the distinction between columnar cell cytoplasm and membrane— minimally differentiated in the raw excitation-emission maps—is visibly improved in DREAM spectra, as shown in the phasor and spectral plots. This suggests that DREAM not only dimensionally encodes spectral information, but also increases spectral separability by concatenating the post-excitation (Stokes-shifted) emission bands across different lasers into a single composite spectrum, yielding larger phasor Angular phase spread (Fig. 3C) and higher CQE (Fig. 5) than single-excitation images. By integrating these peak responses across multiple excitations, DREAM captures a broader and more complementary set of spectral features than is possible with any individual excitation. This cumulative view enhances the separability of biologically distinct regions, such as cytoplasm and membrane, which would otherwise exhibit overlapping or indistinct spectral profiles when analyzed under a single excitation wavelength.

### 3.3 SEER Visualizations Show Enhanced Perceptual Contrast with DREAM

To assess the visual impact of DREAM compression, we applied SEER mapping to generate color-encoded images for each spectral dataset (Fig. 4). All images employ the gradient descent SEER colormap previously validated for autofluorescence phasor analysis [27]. The gradient descent map separates spectra with a bias on subtle spectral differences closer to the center of the phasor histogram. The color palette is fixed and utilizes SEER’s autoscale mode to linearly rescale the colormap for each dataset to maximize the usage of color gamut, ensuring that any perceived difference in contrast arises from the underlying spectral content, not from palette re-tuning. This approach enables direct visual comparison. Fig. 4a displays the DREAM-SEER image, which exhibits strong visual separation among tissue types. Columnar cells are highlighted in red, connective tissue and membranes in distinct tones, and blood cells appear in lighter hues—indicative of a broader spectral distribution. In contrast, single-excitation SEER images (Figs. 4b–4h) show more limited contrast: 405 nm emphasizes nuclei (Fig. 4b), 561 nm highlights membranes (Fig. 4f), and 633 nm selectively enhances blood features (Fig. 4h). Although some single-excitation SEER images accentuate specific structures, none match the overall spectral richness and separability achieved by the DREAM-SEER composite.

### 3.4 Quantitative Assessment of Image Quality

To complement visual inspection, we performed quantitative analysis of SEER image quality using no-reference perceptual metrics: colorfulness, contrast (CRME), sharpness, and the Color Quality Enhancement (CQE) index. All datasets were represented utilizing the same SEER gradient-descent colormap, linearly radially scaled to the dataset’s phasor representation according to SEER approach[27]. These metrics are computed across sliding windows of varying sizes (5 to 45 pixels), to simulate features of different spatial scales. We report results specific to the Fig. 4 dataset (Fig. 5).

For window size 5×5, representing small features such as membranes, the DREAM image achieved the highest median sharpness (44.8), outperforming all single-excitation datasets (e.g., 31.3 for 633 nm). Although colorfulness and contrast (CRME) ranked second-highest and third-highest for DREAM respectively, the combined CQE score was highest (8.48), followed closely by the 405 nm image (8.28). This confirms the visual impression from Fig. 4.

As the window size increased (15×15 and larger), DREAM maintained the highest median sharpness, with values remaining above 50.2 across scales. Notably, colorfulness for DREAM improved at larger scales, surpassing all single-excitation images starting at window size 15 with a median value of 1.29 (e.g., 1.10 for 514 nm). CQE values of DREAM datasets increased accordingly, reaching 9.36 at window size 15, compared to 8.989 for 405 nm. At window sizes 25, 35, and 45, DREAM consistently retained the top CQE score, with sharpness and contrast remaining stable.

Quantitative analysis across a broader set of 19 SEER datasets acquired from 3 different tissue samples further substantiated the performance advantages of DREAM (Fig. 6). Each dataset is evaluated using the same perceptual quality metrics, colorfulness, sharpness, contrast (CRME), and CQE, across five window sizes (5, 15, 25, 35, and 45 pixels) to assess multi-scale consistency. DREAM datasets consistently exhibit the highest colorfulness across all spatial resolutions, with narrow interquartile ranges indicating stable chromatic richness among datasets. This reinforces DREAM’s ability to preserve and enhance spectral diversity across biological specimens.

Sharpness measurements follow a similar trend, with DREAM maintaining the highest median values across all window sizes. At window size 15, corresponding approximately to mid-sized structural elements such as cell nuclei, DREAM achieved a median sharpness above 50, notably outperforming all single-excitation images. The box plots also show reduced variability for DREAM in this metric, underscoring the technique’s reliability in preserving fine tissue structures.

While contrast (CRME) rankings placed DREAM consistently in the top tier, 633 nm excitation achieved the highest contrast values at some window sizes (Fig. 5C). Unlike other excitation wavelengths, where contrast values tend to increase or remain stable with larger window sizes, the 633 nm contrast profile showed an inversion, gradually decreasing as window size increased. This deviation suggests an atypical spatial distribution of contrast in the 633 nm dataset. A plausible explanation is the masking process applied to eliminate specular reflections from laser scatter in the 633 nm excitation, which may have disproportionately affected localized high-intensity regions that dominate small-scale contrast but contribute less at broader spatial resolutions. When aggregating all three perceptual metrics into the composite CQE score, DREAM retained the top rank across all tested spatial scales (p-value<0.02). The CQE distribution for DREAM showed both the highest median and narrowest spread, highlighting its robust performance and cross-sample consistency.

These results confirm that the DREAM-SEER pipeline offers superior perceptual quality in label-free tissue imaging, independent of feature size or dataset variability.

## 4. Discussion

The results presented in this study highlight the effectiveness of DREAM as a robust approach for enhancing visualization of spectral autofluorescence imaging across multiple spatial scales. DREAM works by integrating high-content emission regions from multiple excitation wavelengths into a single 32-channel emission spectrum. It selects the most informative emission segments following each excitation wavelength, combining them into a single, composite spectrum that emphasizes high-content, non-redundant spectral regions. By doing so, DREAM retains the biologically rich and excitation-specific information while reducing spectral redundancy, emphasizing complementary excitation-specific contributions and producing a unified spectrum with both enhanced interpretability and molecular specificity. The result is a compact, high-information dataset that enables better visualization of the biochemical diversity of endogenous tissue autofluorescence.

In particular, DREAM visualization consistently outperforms conventional single-excitation SEER in both qualitative and quantitative assessments of image quality. Metrics such as colorfulness and sharpness, which reflect chromatic richness and preservation of structural detail respectively, are highest for DREAM across nearly all analysis window sizes and, consequently, feature sizes (Fig. 6). These results underscore the benefit of aggregating information from multiple excitation-emission pairs, enabling DREAM to more comprehensively capture the complexity of endogenous tissue fluorescence.

The analysis reveals that the phasor positions associated with the DREAM approach are more widely separated in phasor space compared to single-laser acquisitions (Fig. 3a,b), indicating that the underlying spectra are more distinct across different tissue features. This separation suggests improved spectral discriminability at the pixel and feature level, which directly contributes to enhanced contrast and tissue segmentation. The overlaid spectra for these features (Fig. 3c) further confirm this, showing that DREAM produces more distinct spectral profiles compared to single-laser excitation datasets (Fig. 3d–j). Owing to the underlying calculations, the DREAM spectra do not follow the smooth, asymmetric-bell-shaped profiles typical of isolated fluorescent molecules. Instead, they are piecewise smooth, likely reflecting the contribution of multiple overlapping fluorophores excited across the range of wavelengths. This composite spectral signature may encode richer biological information, enabling more accurate tissue classification.

While certain individual laser excitations (e.g., 514 nm, 561 nm) may highlight some features at a specific scale, DREAM delivers robust image quality across all tested window sizes and spatial scales. The aggregation of multi-excitation emissions in DREAM effectively captures a wider range of spatial and spectral details, enhancing both local and global image quality from a qualitative (Fig. 4) and quantitative (Fig. 5) perspective. The 633 nm excitation achieved the highest contrast scores (Fig. 5c), owing to a combination of biological signal characteristics and a technical factor. The strong optical properties of hemoglobin in blood vessels at this excitation wavelength[35] increase signals in sparse regions of the sample. In addition, the removal of reflectance artifacts from the laser required masking of the dataset, creating local small areas of sharp intensity changes. As a result, DREAM yielded the second-highest average contrast, while maintaining a distinctly narrow distribution across datasets. This convergence of strong contrast and low variance, suggests that DREAM provides both striking and consistently reliable visualization performance across different tissue types and experimental conditions. Consistency is further reflected in the CQE scores: DREAM achieved the highest CQE at every window size tested, indicating its superior overall perceptual quality when colorfulness, sharpness, and contrast are considered together.

An important observation arises from the behavior of the metric distributions at different spatial scales (window sizes). For contrast, DREAM maintained a consistently narrower distribution width, outperforming other excitation wavelengths in terms of uniformity. For colorfulness, sharpness, and CQE, the variance across datasets decreased with increasing window size. Selected window sizes correspond to the various biological feature dimensions found in our dataset. The sizes of the distinguished biological features provide context for these trends: nuclei ranged from ∼35–59 pixels in length, blood regions spanned ∼40–74 pixels, and columnar epithelial cells, including cytoplasm and full cell diameters, reached up to ∼240 pixels in length, with membrane thicknesses as small as ∼17–47 pixels. At window sizes of 35x35 pixels and larger, DREAM not only achieved the highest mean values but also exhibited the narrowest distribution, demonstrating both high fidelity and low variability in representing larger biological features. This trend supports the notion that DREAM produces images with both high visual clarity and consistency, critical for pathology applications where interpretability and reproducibility are paramount.

The results presented here demonstrate clear advantages of the DREAM approach in terms of spectral discrimination and perceptual image quality; however, it must be acknowledged that the dimensionality reduction strategy implemented in this study represents a deliberately simple formulation. By selecting the most informative emission channels following each excitation and concatenating them into a unified spectrum, DREAM achieves a balance between computational efficiency and enhanced contrast. It is likely that more sophisticated dimensionality-reduction or fusion strategies, such as those leveraging machine learning or nonlinear optimization, may further refine performance. We anticipate that such methods would offer incremental gains rather than transformative improvements when benchmarked against the substantial enhancements already observed in DREAM relative to conventional single-laser excitation datasets.

This first demonstration was conducted on esophageal tissue, which includes multiple histological components but does not encompass the full diversity of tissue autofluorescence encountered in other organs. Because DREAM operates directly on the measured excitation– emission data without organ-specific priors, it should, in principle, be applicable to tissues such as brain, liver, or kidney, provided that their dominant intrinsic fluorophores fall within the instrument’s excitation and detection range. Tissues with markedly different biochemical composition may present distinct spectral profiles, and formal cross-tissue validation will be essential to confirm performance in those contexts.

Regarding sample preparation, all specimens imaged here were formalin-fixed paraffin-embedded archival blocks stored 6 years prior to the imaging in this work, as required by institutional regulations for pathology core access. Within this cohort, DREAM’s performance was consistent, suggesting a degree of robustness to long-term storage; however, its sensitivity to other fixation chemistries, preservation conditions, or a broader age range of samples remains untested. Future studies should explicitly evaluate these variables to establish the generalizability of DREAM across tissue types and preparation protocols.

A second consideration concerns the inherent tradeoff in DREAM’s compressive design. By design, DREAM reduces the full excitation-emission map, a high-dimensional spectral representation of tissue autofluorescence, into a lower-dimensional form. While this approach effectively distills salient information across multiple excitations, it also necessarily discards aspects of the joint excitation-emission structure that may encode subtle biochemical or spatial variations. The excitation-emission map likely contains latent features that could be more fully exploited by future computational frameworks capable of multi-dimensional unmixing or fingerprinting. In specialized settings, when a known set of autofluorescent biomarkers is of primary interest, a reduced, task-optimized subset of excitation lines could further improve acquisition speed or spectral contrast, but such tailoring would render the protocol sample-specific rather than broadly deployable. While DREAM provides an effective and practical solution within current imaging pipelines, these findings motivate the development of next-generation algorithms that can operate directly on the full hyperspectral excitation-emission space to unlock additional sensitivity and specificity in biological tissue analysis.

## 5. Conclusions

In this study, we present a powerful dimensionality reduction strategy, Dimensionality Reduction for Enhanced Autofluorescence Microscopy (DREAM), designed to integrate multi-excitation autofluorescence emission data into a single, high-content spectral representation. When applied to hyperspectral datasets of unlabeled esophageal pathology slides, DREAM significantly improves spectral separability between tissue components, offering a path toward more objective and quantitative label-free pathology.

DREAM enhances spectral discrimination while reducing data dimensionality by summarizing key emission features following each excitation and minimizing redundant or low-information spectral channels. Coupled with phasor-based SEER visualization, this approach enables intuitive yet quantitative color mapping of autofluorescent tissue without the need for exogenous labels or chemical stains. Our results show that DREAM-SEER images consistently outperform single-excitation SEER in both visual and quantitative assessments, including colorfulness, contrast, sharpness, and the integrated CQE score. This improvement is robust across multiple spatial scales, underscoring the method’s ability to preserve biologically relevant features of varying size.

Although DREAM was developed and validated here in the context of label-free pathology on fixed tissue sections, the underlying principles are broadly generalizable to any multi-excitation fluorescence imaging modality. Importantly, the excitation-emission data used here reflect intrinsic signals from endogenous fluorophores, and the success of DREAM in resolving complex tissue architecture without stains reinforces the potential of autofluorescence-based diagnostics. The dimensionality reduction implemented in DREAM represents an efficient first step. It remains possible the full excitation-emission datasets contain additional information that could be exploited by more sophisticated multidimensional unmixing algorithms. Future work may focus on developing such models to further harness the diagnostic power of hyperspectral autofluorescence imaging.

## 6. Acknowledgement

The authors thank C.-L. Chiu (Chan Zuckerberg Initiative) for helpful discussions. This material is based upon work supported by the Translational Imaging Center and Bridge Institute at University of Southern California.

## 7. Data Availability

Data underlying the results presented in this paper are available upon reasonable request. An example dataset (Figure 4) is available on our website https://bioimaging.usc.edu/software.html

